# GeneSIS: enhancing transferability of polygenic scores with variant-level gene-by-sex interaction effects

**DOI:** 10.64898/2026.09.14.751452

**Authors:** Yosuke Tanigawa, Manolis Kellis

**Affiliations:** Computer Science and Artificial Intelligence Laboratory, Massachusetts Institute of Technology, Cambridge, MA, USA; Broad Institute of MIT and Harvard, Cambridge, MA, USA; Department of Bioengineering, University of California, Los Angeles, Los Angeles, CA, USA

## Abstract

Advancing precision medicine requires accurate prediction of disease liability across populations and contexts. A major challenge is the limited transferability of polygenic scores (PGS) across genetic ancestry groups. We present GeneSIS (GENE and Sex Interaction Score), a supervised statistical learning framework for jointly modeling additive and context-dependent genetic effects at single-variant resolution directly from individual-level data. We analyze 406,659 individuals, including admixed individuals, in the UK Biobank and 1.3 million variants to develop predictive models for 99 complex traits. We report that ∼8% of selected variables capture gene-by-sex (GxS) effects, validated by sex-stratified analyses. Modeling GxS effects improves prediction across 32 traits in non-European individuals. For predicting hip circumference in Africans, GeneSIS achieves a 3.7-fold improvement (p=8.0×10-7) over linear-only PGS and highlights biologically plausible hypotheses, such as pleiotropic GxS effects of *GCKR* (rs1260326) on anthropometry and menopause, as well as GxS pathway enrichments for interleukin-4 regulation. Overall, our results highlight the benefits of integrating context-dependent effects in human genetics studies.

## Introduction

Predicting disease liability and medically relevant traits from individuals’ genetic, demographic, and environmental factors has substantial implications in epidemiology, medical prevention and intervention strategies, and, more broadly, in realizing precision medicine. Conventional risk factors, for example, age, biological sex, family history, and smoking status, have been extensively studied and used in clinical practice. The incremental utilities of genetic factors have recently attracted substantial research interest[1].

Polygenic score (PGS) is a statistical approach to aggregate genetic effects across multiple genetic variants into a single score per individual[1]. Recently developed PGS models show improved predictive performance, highlighting their potential clinical relevance for some traits[1–4]. The predictive performance of PGS shows substantial variability across genetic ancestry groups, posing potential concerns about their applications[5]. Despite growing recognition of shared causal genetic effects across genetic ancestry groups[6], PGS models trained on a single population do not translate well to other population groups. The limited predictive performance is partly due to differences in linkage disequilibrium structure and allele frequencies across continental ancestries and, in some cases, ancestry-dependent genetic effects. Several efforts are underway to better characterize genetic effects across diverse contexts and translate findings into better predictive models, including active recruitment of ancestry-diverse individuals in genetic studies and computational method development [6–9]. Emerging computational approaches integrate genetic data from multiple ancestry groups or individuals across the continuum of genetic ancestry, as exemplified in our recently proposed inclusive PGS model[9]. PGS can be combined with other factors, such as family history, conventional risk factors, and rare variant genetic effects[10]. Due to technical limitations, most PGS approaches do not account for nonlinear (non-additive) or context-dependent genetic effects. They are mostly limited to linear effects captured in summary statistics from genome-wide association studies (GWAS)[8,12], even though most common complex traits are influenced by a combination of conventional risk factors, demography, genetic components, and their interactions.

Recent studies highlight the substantial roles of nonlinear and context-dependent genetic effects on complex traits, thanks to larger sample sizes and methodological advances [11–21]. On the one hand, nonlinear genetic effects, such as genetic dominance and gene-by-gene (GxG) interactions (also known as epistasis), contribute smaller but substantial amounts to heritability than linear effects[14,15]. On the other hand, studies on context-dependent genetic effects reveal substantial gene-by-sex (GxS) and gene-by-environment (GxE) interactions[16–19,22–27]. For example, substantial GxS effects have been reported for testosterone, anthropometry, and blood pressure traits[17,18,22–26,28,29]. Genetic effects on molecular traits (e.g., expression quantitative trait loci effects) are modulated by cellular contexts, including cellular states, stimulus-response, and exposure to drugs[13,21]. Examples of studying context-dependent genetic effects also manifest when mapping ancestry-specific or ancestry-dependent genetic effects on heritable traits. Differences in the environment and the resulting GxE interactions in the present and the past, and population bottleneck events, have contributed to shaping ancestry-specific or ancestry-enriched alleles, some of which show associations with complex traits[30–32]. A recent study has proposed several distinct explanations underlying context-dependent genetic effects, including locus-specific modulation of genetic effects, varying genetic variance, and proportional amplification of both genetic and environmental variance[27]. Under the first scenario, the genetic correlation between different contexts would be smaller than 1, indicating that the genetic association summary statistics characterized under an additive model may not fully represent the complexity of genetic effects across environmental contexts. These highlight the potential advantage of integrating linear, nonlinear, and context-dependent genetic effects in a unified framework.

Indeed, a few pioneering works by us and others have started to incorporate nonlinear and context-dependent effects into predictive models, one at a time[9,23,24,33–35]. For nonlinear effects, for example, we recently developed GenoBoost, a flexible machine-learning-based PGS modeling framework, to incorporate nonlinear genetic dominance effects and showed that genetic dominance effects localized in the major histocompatibility complex (MHC) locus improve predictive accuracy for immune-related disorders[33]. For gene-by-demography interactions, we showed that considering interactions between genotype PCs and genetic variant effects at a relatively small number of genetic loci can improve the accuracy of genetic prediction[9]. For GxS interaction effects, we revealed that constructing sex-stratified PGS models for testosterone, which shows an exceptionally high degree of sexual dimorphism in its genetic architecture, improved the predictive performance despite the two-fold reduction in the sample size in the PGS training population[23]. Beyond testosterone, there has been limited success in improving the predictive performance of PGS by incorporating GxS or GxE interaction effects[17,25,36]. Nonetheless, applying sex-stratified PGS analysis highlights its utility in accurately predicting age-dependent temporal dynamics of hematological traits[34], and it has been proposed that amplification (e.g., varying magnitude of the causal effects between male and female) plays a substantial role in shaping GxS interaction effects[24]. A recent study proposes a calibration of PGS to account for context-dependent prediction intervals of PGS models[35]. However, prior approaches are limited because they often assess the nuanced genetic effects in aggregate across multiple genetic variants, require stratification of samples (e.g., male- and female-only cohorts in sex-stratified analysis), or focus on a specific type of nonlinear or context-dependent effects at a time.

Here, we overcome these limitations and present GeneSIS (GENE and Sex Interaction Score), a unified predictive modeling framework capable of integrating linear and nonlinear effects of millions of genetic variants and their context-dependent effects. We hypothesize that incorporating nonlinear and context-dependent effects can better capture nuanced effects and help improve the transferability of PGS. To that end, GeneSIS substantially extends the recently developed inclusive PGS (iPGS)[9]. GeneSIS and iPGS fit predictive models directly on individual-level data, thus naturally applicable to individuals across the continuum of genetic ancestry. As an initial application of GeneSIS, we analyze n=406,659 unrelated, ancestry-diverse individuals in the UK Biobank and develop predictive models for 99 quantitative traits, considering linear and nonlinear effects of genetic variants and GxS interaction effects for each genome-wide genetic variant[37]. We validate our approach using sex-stratified GWAS analysis and report substantial improvements in PGS transferability for non-European individuals. We report significant gains in predictive performance for individuals of African ancestry, most notably in anthropometric traits, including body mass index and hip circumference, which are relevant measures of obesity. Furthermore, we show that GeneSIS reveals biologically plausible hypotheses. For example, we show that genetic variants with GxS effects have pleiotropic associations with sex-specific factors, and genome-wide GxS effects for hip circumference are enriched for known relevant biological processes in obesity, nominating attractive therapeutic targets for context-dependent intervention.

## Results

### GENe and Sex Interaction Score (GeneSIS) methodology

In GeneSIS, we consider linear and interaction effects of genetic variants, demography, and biological sex by applying supervised learning. In this initial study, we used the following steps as a proof of principle, though a broader set of approaches is possible, as described in the discussion. Here, we first augmented genetic data by constructing variables representing nonlinear genetic effects (**Fig 1A, Methods**). Second, we applied supervised statistical learning directly on the individual-level data while introducing more regularization for nonlinear and context-dependent genetic effects. Third, we evaluated the incremental utility of nonlinear and context-dependent genetic effects in polygenic prediction by benchmarking GeneSIS against linear-only inclusive PGS (iPGS) models. As an illustration of follow-up analysis, we nominate biologically relevant variants, genes, and enriched pathways captured in the GeneSIS predictive model.

**Fig 1.**
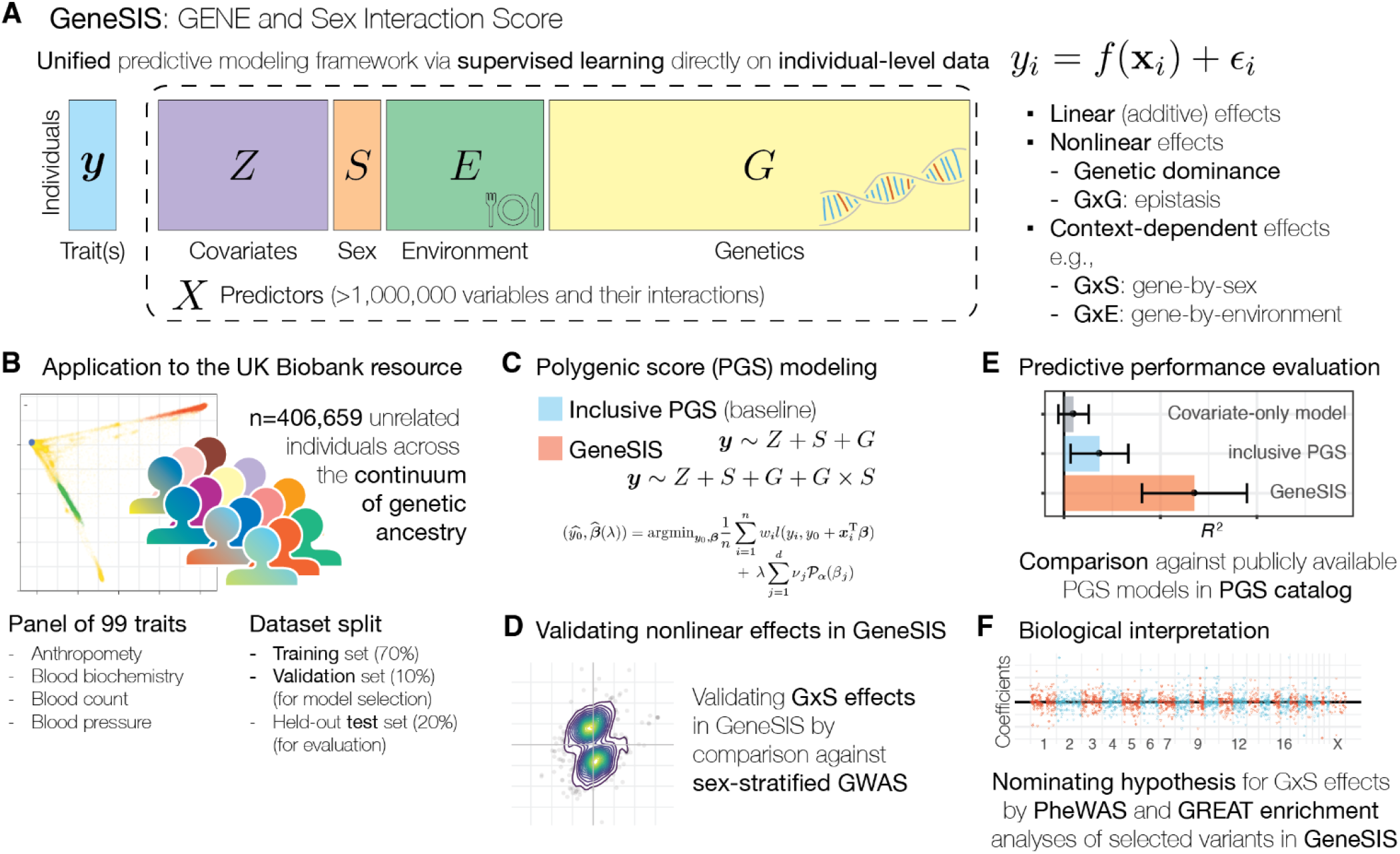
Illustrative overview of the study. (**A**) GeneSIS is a unified predictive modeling framework using supervised learning directly on individual-level data. GeneSIS considers linear (additive), nonlinear, and context-dependent genetic effects. (**B**) Initial application of GeneSIS focusing on variant-level genome-wide GxS effect to 99 quantitative traits in the UK Biobank resource across n=406,659 ancestry-diverse individuals. (**C**) Polygenic score (PGS) modeling with GeneSIS and inclusive PGS (iPGS). Inclusive PGS is a baseline method capable of considering linear effects alone. (**D**) Validation of nonlinear effects captured in the GeneSIS model. (**E**) Predictive performance evaluation of GeneSIS. (**F**) Biological interpretation of genetic variants captured in the GeneSIS model.

### Overview of application of GeneSIS to UK Biobank

In our application of GeneSIS to n=406,659 unrelated individuals across the continuum of genetic ancestry in the UK Biobank resource, we assembled a panel of 99 heritable traits, consisting of anthropometry, blood biomarkers, blood pressure, and hematological traits (**Fig 1B, Table S1-S2**)[37]. We applied *L*_1_- and *L*_2_-penalized Elastic Net regression directly on the individual-level data considering linear, nonlinear, and genome-wide gene-by-sex (GxS) interaction effects, represented in 2,630,335 predictor variables across 1,316,147 genetic variants (**Fig 1C, Methods, Table S3**)[9,38–41]. For the MHC region, we used the imputed human leukocyte antigen (HLA) allelotypes to account for complex linkage disequilibrium (LD) structure and also incorporated genetic dominance effects[33]. We applied linear-only iPGS models as a baseline using the same set of 1.3 million genetic variants[9]. In both GeneSIS and iPGS models, we used an additional 23 variables as unpenalized covariates, including age, sex, age^2^, age*sex, Townsend deprivation index, and the first 18 genotype PCs. We split each population group and used the training set for fitting models, the validation set for hyperparameter tuning and model selection, and the held-out test set for predictive performance evaluation (**Table S1**).

Of the analyzed individuals, about 53.9% are female (**Table S1**). The unpenalized covariate term, sex, accounts for the mean differences in trait values between male and female individuals. To allow the magnitude or direction of genetic effects to differ between male and female groups and to capture nuanced genetic effects, we prepared about 50.0% of predictor variables to represent GxS interaction effects (**Table S3**).

We first demonstrated that GeneSIS’s GxS effects are highly consistent with sex-stratified genome-wide associations, validating our approach (**Fig 1D**). We subsequently evaluated the predictive performance of GeneSIS, linear-only iPGS, and a model that considers covariate terms alone in each of the following population groups: White British (WB), non-British white (NBW), South Asian (SA), African (Afr), and other unrelated individuals (Others) (**Fig 1E**). Lastly, we highlight the genetic variants selected in the GeneSIS model and their enrichment to pathways and biological processes[42,43] to offer an interpretation of context-dependent effects captured in the predictive model (**Fig 1F**).

### Validating GxS effects in GeneSIS by sex-stratified GWAS

Across the 99 quantitative traits, we developed the GeneSIS predictive model. We also prepared linear-only iPGS models using the same sets of individuals and genetic variants[9]. Both approaches use penalized regression (Elastic Net) directly on the individual-level data without requiring users to specify the genetic architecture of the trait. As such, our study design is suitable for systematic comparison across a wide variety of traits, enabling systematic assessment of the benefits of incorporating nonlinear and context-dependent effects in genetic prediction.

Overall, we found that GeneSIS and linear-only iPGS models have a comparable number of predictor variables (linear regression fit of y=1.006x, Pearson’s correlation=0.97, 95% confidence intervals, CI: [0.96, 0.98]), except for total bilirubin and direct bilirubin (**Fig 2A, Table S4**). The number of predictor variables increased from 197 to 14,869 for total bilirubin and from 278 to 9979 for direct bilirubin in GeneSIS. In GeneSIS models, a median of 8.0% of predictors (ranging from 0.2% to 27.7%) capture GxS interaction effects. Variants with GxS interaction effects are distributed genome-wide, not necessarily localized to sex chromosomes. On a median across 99 UK Biobank traits, the GxS interaction effect size is 24.8% smaller than that of linear effects (**Fig S1**). Genetic dominance effects at HLA allelotypes are captured in two traits (gamma-glutamyl transferase and C-reactive protein), both on the same allele, HLA-DPA1*0103. The high allele frequency of the allelotype (96.0% in White British) works advantageously in capturing genetic dominance effects, possibly reflecting limited statistical power to detect and capture genetic dominance effects for other imputed allelotypes.

**Fig 2.**
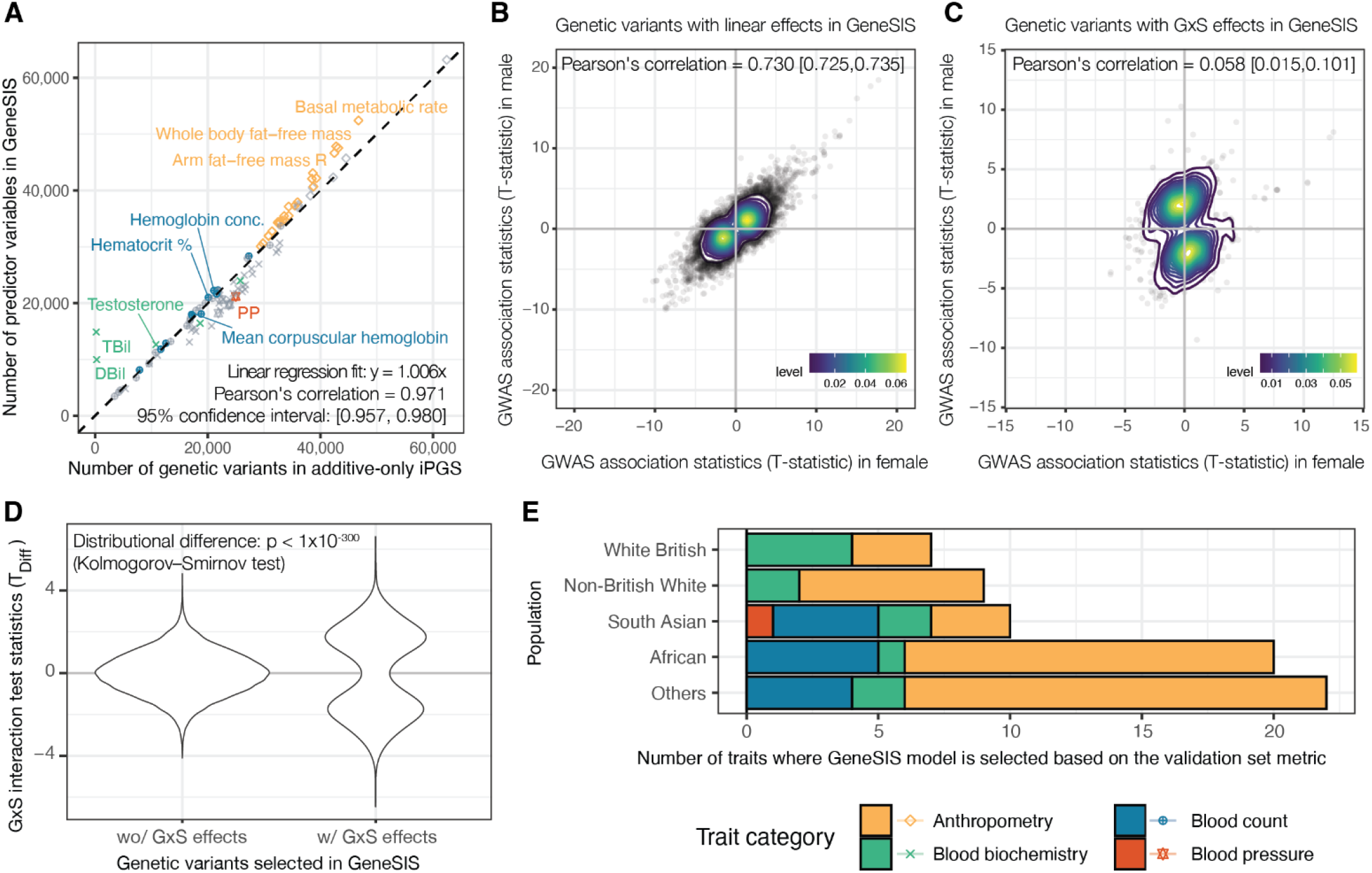
GeneSIS integrates validated GxS effects on top of linear effects in a unified framework. (**A**) The number of predictor variables considered in iPGS (x-axis) and the GeneSIS model (y-axis). Each point is colored by the trait category and whether the GeneSIS model is selected based on the validation set metric. TBill: total bilirubin. DBill: direct bilirubin. (**B**-**C**) Comparison of GWAS effect size estimates between females (x-axis) and males (y-axis) for hip circumference in White British individuals. The genetic variants selected in GeneSIS only for linear effects are shown in (**B**), and the selected variants with GxS interaction effects are shown in (**C**). (**D**) The distributions of GxS interaction effect test statistics, *T*_Diff_, for hip circumference in White British individuals. We show genetic variants selected in the GeneSIS model, stratified by whether the variant has GxS interaction effects. (**E**) The number of traits where GeneSIS is selected for trait prediction on the basis of the validation set metric. For each population group (y-axis), the number of traits per trait category is shown.

To validate the GxS interaction effects captured in GeneSIS models, we applied sex-stratified GWAS analysis and independently validated the presence of GxS effects. To that end, we focused on unrelated individuals in White British, the largest population group in the UK Biobank cohort, and applied sex-stratified GWAS for quantile-normalized phenotypes (**Methods**). We compared effect size estimates and assessed the heterogeneity of genetic associations between male and female individuals. When we investigated the genetic variants selected in GeneSIS, we found that the GWAS association summary statistics from the sex-stratified analysis are more concordant in variants selected for linear effects alone (**Fig 2B-D**). For hip circumference, for example, Pearson’s correlations of association summary statistics were 0.730 (95% confidence interval, CI: [0.725, 0.735]) for genetic variants with linear effects in GeneSIS and 0.058 (95% CI: [0.015, 0.101]) for genetic variants with GxS interaction effects (**Fig 2B-C**). We further quantified the magnitude of GxS effects by computing the variant-level test statistic, *T*_Diff_, where deviation from *T*_Diff_ = 0 indicates the presence of sex-biased genetic effects (**Methods**)[18]. Indeed, the distribution of the *T*_Diff_ statistics is different between the genetic variant groups with linear-only and GxS effects (Kolmogorov–Smirnov test p-value < 1 × 10^−300^) (**Fig 2D**). A similar pattern was observed for other traits, including the two bilirubin traits (**Fig S2**). Together, those results highlight the unique ability of GeneSIS to capture validated GxS effects in predictive models.

### GeneSIS model selection identifies trait- and population-specific benefits

Having validated the GxS effects, we evaluated the predictive performance of GeneSIS models and compared them against the corresponding linear-only iPGS models. We report predictive performance in each population group to minimize the risk of confounding due to population structure (**Table S5**). We subsequently tested the statistical difference between the two models in the validation and held-out test sets (**Table S6**). When comparing GeneSIS and iPGS, we focused on full models that include covariates, genetics, and their interactions, and evaluated their predictive performance.

Across 99 traits, we found substantial heterogeneity in improvements in predictive performance with GeneSIS, consistent with differences in GxS genetic architecture across traits. We applied model selection between GeneSIS and linear-only iPGS separately for each (population, trait) pair using population-matched validation sets to reflect target-aware deployment scenarios (**Methods, Fig S3**). We found that GeneSIS was selected for 41 out of 99 traits for at least one population group (**Fig 2E, Table S7**). The frequency of GeneSIS selection varied across populations, ranging from 7 of 99 traits (7.1%) in White British individuals to 20 of 99 traits (20.2%) in African individuals and 22 of 99 traits (22.2%) in the Others group (**Table S8**). For example, in African individuals, GeneSIS was selected for 14 of the 31 anthropometric traits considered in the study.

We then evaluated whether model selection based on the validation set generalized to the held-out test set. Among the 68 (population, trait) pairs for which GeneSIS was selected, GeneSIS showed equivalent or superior predictive performance to linear-only iPGS in the held-out test set for 66 pairs (97.1%; **Table S9**). The remaining two pairs were observed in the non-British White population. For those two cases, GeneSIS and the linear-only iPGS showed nearly equivalent predictive performance, with *R*^2^ differences of less than 0.0017 in the held-out test set. These results indicate that validation-based model selection effectively identifies settings in which incorporating GxS effects improves or maintains predictive performance in held-out individuals.

### GeneSIS improves PGS transferability, especially for anthropometric traits

Empirically, we found improved predictive performance of GeneSIS models for non-European population groups, most notably in African individuals and for anthropometric traits, including hip circumference, body mass index, and whole body fat mass (**Fig 3A-B**). The magnitude of improvement varied substantially across traits and was not directly correlated with the fraction of predictor variables with GxS interaction effects (**Fig S4**-**S5**). Nonetheless, GeneSIS was selected for many anthropometric traits, consistent with a recent study reporting locus-level gene-by-sex interactions for anthropometric traits[27].

**Fig 3.**
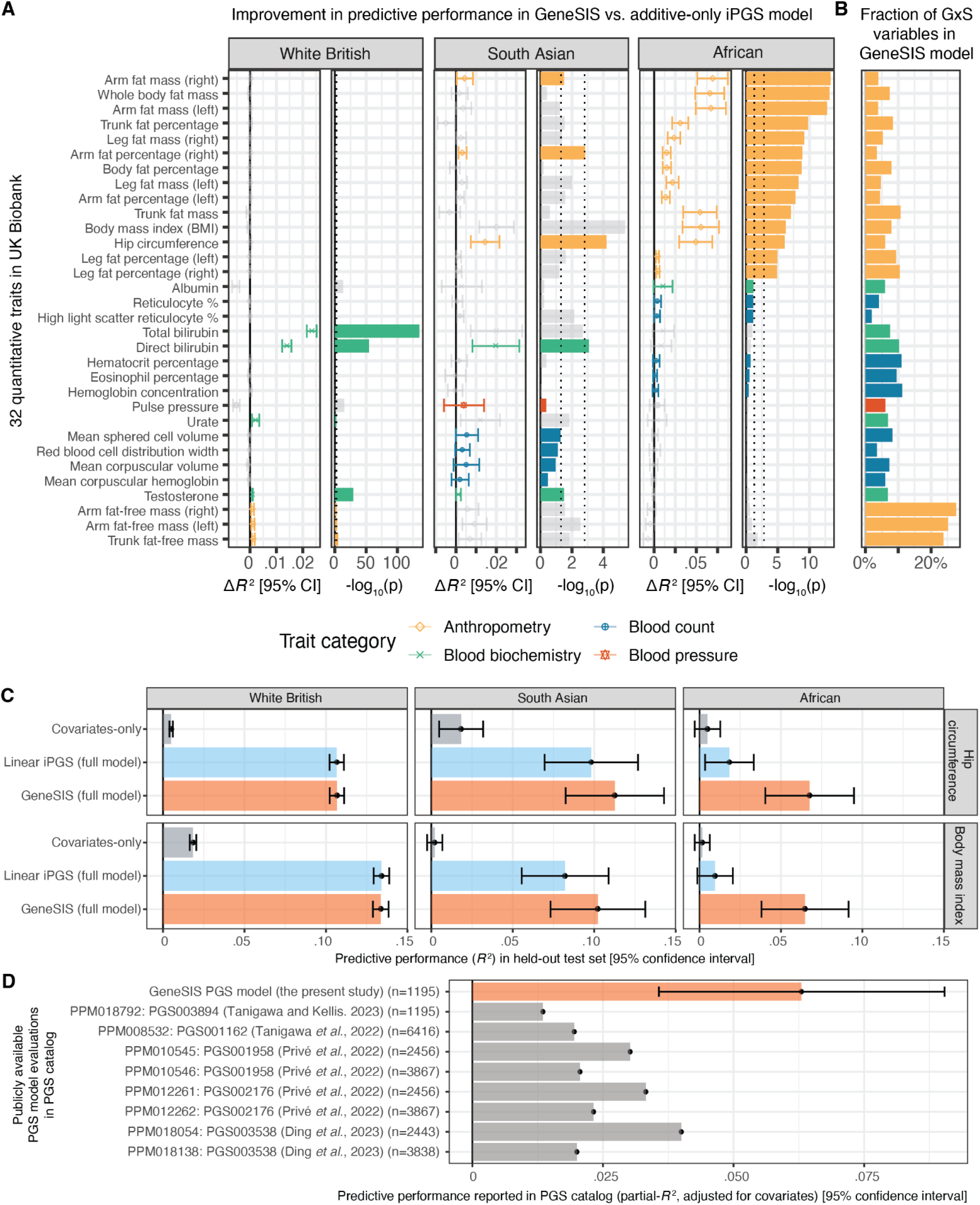
Enhanced PGS transferability in non-European populations with GeneSIS. (**A**) The magnitude and statistical significance of the gain in predictive performance of GeneSIS over linear-only iPGS across three population groups. We show 32 traits where GeneSIS is selected for at least one of the three populations (White British, South Asian, and African). We show the full results in **Fig S4**. We selected the GeneSIS or linear-only iPGS model based on the validation-set metric. The (trait, population) pairs selected by the validation-set metric are shown in color. (**B**) Fraction of GxS interaction terms in the selected predictor variables in the GeneSIS model. (**C**) Predictive performance (*R*^2^) of covariate-only model, linear-only iPGS model, and GeneSIS model for hip circumferences and body mass index in individuals of White British, South Asian, and African ancestry in the held-out test set. (**D**) For hip circumference, we compare the predictive performance of the GeneSIS model (partial *R*^2^ for covariate-adjusted phenotype, x-axis) for individuals of African ancestry against all publicly available PGS model evaluations in the PGS catalog (y-axis). Error bars represent 95% confidence intervals.

For hip circumference in African ancestry individuals, GeneSIS significantly improved prediction in the held-out test set (*R*^2^=0.067, 95% confidence interval: [0.040, 0.095]) compared with iPGS (*R*^2^=0.018, 95% CI: [0.003, 0.033]) with a p-value of 8.0 × 10^−7^ for the difference in predictive performance (**Fig 3C, Fig S6**). We found similar improvements for body mass index in African individuals (**Fig 3C**).

Moreover, GeneSIS outperformed all publicly available PGS model evaluations for hip circumference in African ancestry individuals[9,44–47] (**Methods, Fig 3D, Table S10**-**S11**). Notably, individuals of African ancestry comprise only 1.49% of the GeneSIS training population, and only 6% of the selected predictor variables (2,075/34,351 variables) capture GxS interaction effects.

For hip circumference, we found improved predictive performance in South Asian individuals with GeneSIS (*R*^2^=0.113) compared with iPGS (*R*^2^=0.098), whereas predictive performance in White British individuals remained nearly identical (*R*^2^=0.1067 for GeneSIS and *R*^2^=0.1066 for iPGS). The difference in the White British group was not significant (nominal p-value=0.62). Overall, these results highlight the benefits of incorporating context-dependent genetic effects for improving PGS transferability.

### GeneSIS prediction is robust to alternative encoding of sex-specific effects

To investigate the effects of encoding sex-specific effects in GeneSIS, we selected seven phenotypes, fit additional GeneSIS models, and assessed changes in predictive performance. In the primary analysis, we considered 2.6 million predictor variables across 1.3 million unique genetic variants, including sex-shared and male-specific GxS interaction effects, given that the analyzed UK Biobank cohorts consisted of 53.9% female and 46.1% male individuals and that collinearity in sex-specific GxS effect vectors introduces numerical stability challenges. In the additional analysis, we considered sex-shared, male-specific, and female-specific effects and compared predictive performance (**Methods**). We found that the two models showed highly consistent predictive performance across five population groups, supporting the robustness of the male-specific GxS encoding used in the primary analysis (**Fig S7**).

### Sparse GeneSIS models offer biological interpretations

To investigate the biology of GxS effects captured in the GeneSIS models, we hypothesized that the genetic variants captured in GeneSIS with linear and GxS effects offer interpretation. We found that the GeneSIS model captures genetic effects on non-synonymous coding variants in well-known genes, allowing interpretation of the biological processes captured in the predictive model. We also inferred the biological basis of sex-dependent genetic effects in the GeneSIS model by investigating pleiotropic associations and genome-wide ontology enrichment of genetic variants with GxS effects.

As an illustrative example, here we focused on the GeneSIS model for hip circumference. For linear effects, we found that protein-altering variants in *MC4R* (rs2229616) and *GIPR* (rs1800437) were selected for their trait-lowering effects (**Fig 4A, S8a, S8b**). The melanocortin 4 receptor, encoded in the *MC4R* gene, is known for its role in regulating body weight and energy homeostasis, whereas the gastric inhibitory polypeptide receptor, encoded in *GIPR*, is known for glucose metabolism; both are known for their relevance in anthropometric traits[48,49]. For GxS interaction effects, we found a non-synonymous variant in *GCKR* (rs1260326), where the encoded glucokinase regulator plays a regulatory role in glucose metabolism and is associated with central fat accumulation (**Fig 4B, S8c**)[50]. Moreover, the variant shows pleiotropic associations across blood biochemistry (for example, triglycerides, C-reactive protein, and sex hormone-binding globulin [SHBG] levels), anthropometric (whole-body water mass and trunk fat-free mass), blood pressure (pulse rate and position on pulse wave notch), and sex-specific traits (**Fig 4C, Table S12**)[51–54]. The associations with the sex-specific traits include genetic effects on age at menopause and “had menopause,” a binary questionnaire-based phenotype. The pleiotropic association of the variant across SHBG and sex-specific traits offers insights into the biology behind the sex-biased effects represented in the GeneSIS model.

**Fig 4.**
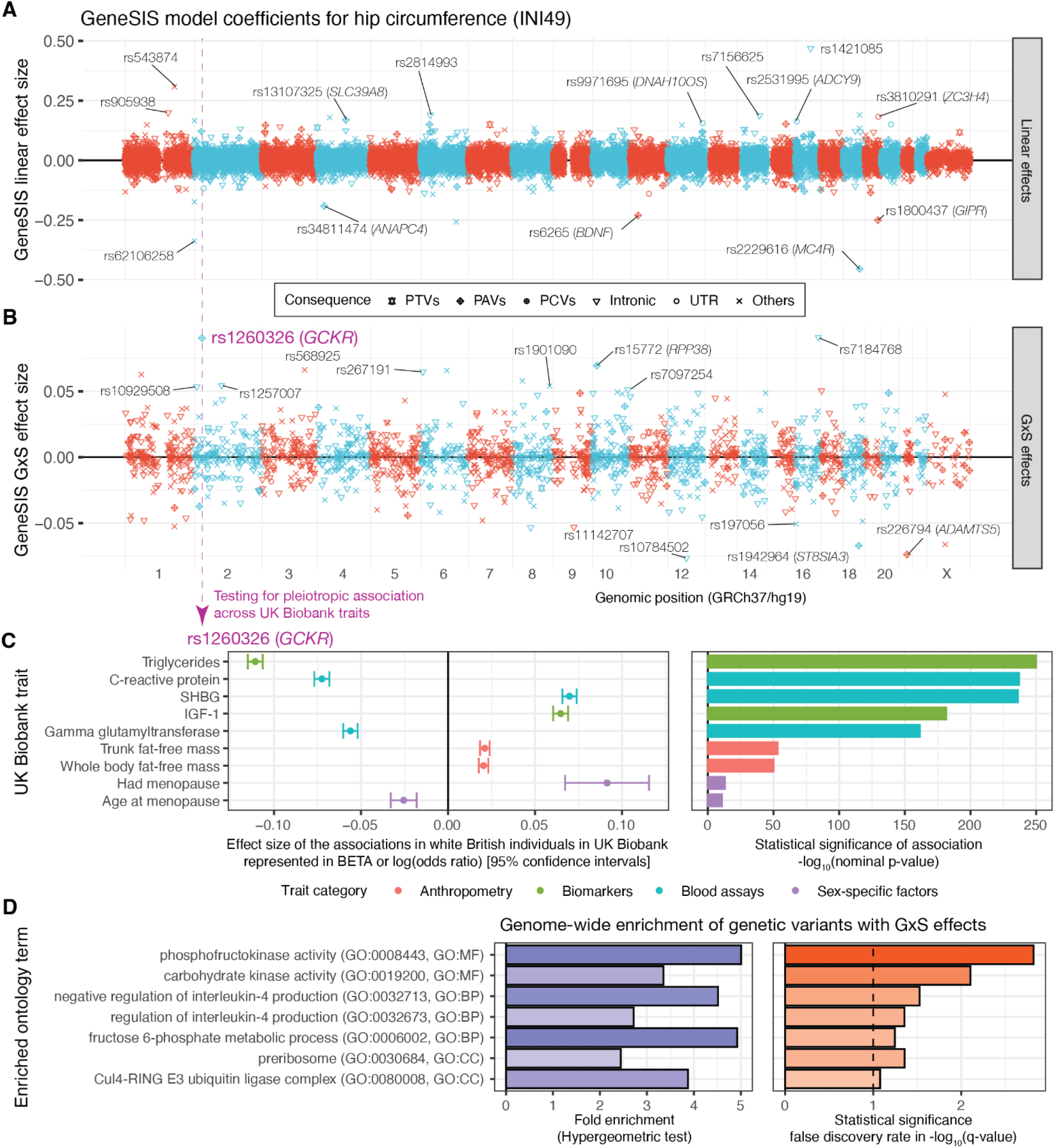
Sparse GeneSIS models offer interpretation. (**A**-**B**) The linear (**A**) and GxS interaction (**B**) effects in the GeneSIS model for hip circumference. We annotated genetic variants with large effects. The GxS effect size directions are represented for male individuals. (**C**) Pleiotropic association of rs1260326, a protein-altering variant in the *GCKR* gene, for select traits. We show the GWAS-based effect size estimates on the left and the statistical significance of the association on the right. Phenome-wide associations with nominal p<1 × 10^−10^ are shown in **Table S12**. (**D**) We show enriched ontology terms for genetic variants with GxS effects in GeneSIS, nominated by the GREAT enrichment analysis.

To further evaluate the biological processes captured in the genome-wide GxS effects in GeneSIS beyond the single-locus analysis, we evaluated the enrichment of ontology terms, pathways, and biological processes using the Genomic Regions Enrichment of Annotations Tool (GREAT) (**Methods**)[42,43]. For hip circumference, we found that the genetic variants selected for GxS interaction effects are enriched for negative regulation of interleukin-4 production (hypergeometric fold change=4.5, FDR=3.0 × 10^−6^) and phosphofructokinase activity (hypergeometric fold change=5.0, FDR=1.51 × 10^−3^) (**Fig 4D, Table S13**). The *PFKP* gene encodes phosphofructokinase, a key enzyme in glycolysis, and Interleukin-4 has been reported to inhibit adipogenesis, regulate lipid metabolism by promoting lipolysis, and influence obesity and type 2 diabetes mellitus [55]. Overall, our results indicate that analysis of genetic variants in GeneSIS reveals key pathways with sex-dependent effects, nominating attractive targets for follow up studies.

## Discussion

Here, we present GeneSIS, a unified polygenic score (PGS) modeling framework capable of incorporating linear, nonlinear, and context-dependent effects. We show its advantage in incorporating nuanced effects in genetic prediction and illustrate the biological characterization of GeneSIS models. As an initial application, we analyze 99 quantitative traits in UK Biobank using n=406,659 unrelated individuals across the continuum of genetic ancestry. First, we validate the gene-by-sex (GxS) interaction effects captured in GeneSIS by an orthogonal GWAS-based approach. We then compare GeneSIS’s predictive performance with linear-only inclusive polygenic scores (iPGS) trained on the same ancestry-diverse cohort. Given substantial heterogeneity across traits and populations, we applied validation-based model selection between GeneSIS and linear-only iPGS, with 66 of 68 GeneSIS selections showing equivalent or superior predictive performance in the held-out test set. We show that modeling GxS effects improves the transferability of polygenic scores, especially for anthropometric traits in African individuals. Of note, the GeneSIS model for hip circumference, an illustrative example, outperforms all publicly available model evaluations for Africans in the PGS catalog.

Lastly, we nominate biologically plausible hypotheses for context-dependent effects captured in GeneSIS. For example, we report phenome-wide associations across menopausal age and sex hormone-binding globulin levels on genetic variants with GxS interaction effects in GeneSIS. Moreover, we find genome-wide enrichment of GxS effects in GeneSIS for biological processes and pathways, nominating attractive targets for follow up studies and potential context-dependent interventions.

Improving the accuracy of predictive models for disease outcomes and medically relevant traits has direct implications for realizing precision medicine. To that end, substantial progress has been made in improving the transferability of PGS. Existing approaches include prioritizing variants present in diverse populations[58] or overlapping with bio-sample-specific regulatory elements[59], integrating statistical fine-mapping results [56], and incorporating genetic data from ancestry-diverse individuals[7,9,56,57]. Here, we reveal that joint modeling of linear, nonlinear, and context-dependent effects, including GxS interactions, is another effective strategy for developing more transferable PGS models, complementing existing efforts. The GeneSIS model does not assume genetic effects are linear or universally shared and allows contextual variables to modulate them. Our flexible approach captured context-dependent effects not represented by conventional additive models and improved PGS transferability across genetic ancestry groups. In our initial application, we observed improvements in predictive performance, mostly for anthropometric traits in individuals of African ancestry. With GeneSIS, the gap in predictive performance between the European and African populations has decreased.

However, predictive performance remains highest in the European population group, reflecting that most participants used in the PGS training are of European genetic ancestry. We envision that insights from the study would be instrumental for further follow-up studies considering gene-by-environment (GxE) interaction effects beyond binarized biological sex and other traits[26,58,59]. Further analyses would reveal when explicitly modeling context-dependent non-linear effects would be beneficial.

Methodologically, our study highlights substantial advancements in the predictive modeling of complex traits. GeneSIS is capable of incorporating linear, nonlinear, and context-dependent genetic effects across millions of genetic variants in a unified framework. We modeled genome-wide linear effects, GxS interactions at single-variant resolution, and genetic dominance effects in the HLA region in a unified framework; we plan to extend this to genome-wide nonlinear and GxE effects. The ability to characterize context-dependent effects at each genomic locus directly from individual-level data allows incorporation of context-dependent effects even in the presence of imperfect genetic correlation across demographic and environmental factors[27]. Here, we focus on penalized regression as an initial application given its success in PGS modeling[9,38,39,45], but other predictive models, including statistical boosting[33,60,61], would be attractive alternatives. The proposed GeneSIS framework is particularly well-suited for capturing locus-specific context-dependent genetic effects, where the direction or magnitude of the genetic effect differs between contexts, such as between sexes, at specific genomic loci. Other forms of gene-by-context interactions, such as varying heritability or proportional amplification of genetic and environmental variance across different contexts, as discussed in a recent publication[27], may be better captured by alternative approaches, including transformation and normalization of phenotypic variables across different contexts or distributional regression techniques[62]. Our results, together with our previous PGS modeling efforts [9,33], highlight that applying supervised learning directly to large-scale individual-level data is an effective strategy for incorporating genetic effects not captured in linear association summary statistics from GWAS studies.

Biologically, we illustrate that sparse GeneSIS predictive models readily offer interpretations. We show that pleiotropic associations with sex-specific factors offer plausible hypotheses on sex-dependent genetic effects captured in the GeneSIS models. We nominate biological processes in obesity for hip circumferences via genome-wide enrichment of GxS effects, offering attractive targets for context-dependent interventions. Further follow-up studies would be helpful, as there is no guarantee that the genetic variants selected in the GeneSIS models have causal roles, as is typical for predictive models. To that end, fine-mapping analysis capable of considering linear, nonlinear, and context-dependent effects would help boost confidence in the mechanisms of action. Increased knowledge of context-dependent genetic effects may also offer insights into the “missing regulation” problem[63].

Our analysis opens several directions for future studies not considered in our initial application presented here. First, we primarily focused on genome-wide GxS interactions and genetic dominance effects at the HLA locus, where we considered imputed allelotypes to account for complex linkage structures in the region, while keeping genome-wide nonlinear encodings as natural extensions. The flexible GeneSIS modeling framework can also capture other nonlinear effects, such as gene-by-gene (GxG) interactions (epistasis) and GxE interactions. Future studies should expand the analysis into different contexts. Applications of variance quantitative trait loci and Mendelian Randomization analyses can nominate relevant genetic variants and environmental factors and potentially reduce the search space[16]. Second, we prioritized linear genetic effects over nonlinear effects and coding variants over non-coding variants, both using penalty factors (**Methods**). In this initial study, we determined the penalty-factor values using heuristics and applied the same values across all 99 traits. In principle, we could optimize hyperparameter values for each trait by considering the trait’s genetic architecture, such as the proportion of heritability explained by linear and GxS interaction effects and heritability enrichment over functional genomic annotations. Emerging regression techniques, such as regression models that consider “feature of features”[64], will be relevant moving forward. Third, we currently focus on the UK Biobank only, given the challenges of sharing individual-level data across cohorts, but future studies will benefit greatly from combining multiple cohorts. Further expansion of our approach may allow joint modeling of linear, nonlinear, and context-dependent effects based on GWAS summary statistics under various context-dependent regression models across multiple traits, cohorts, and contexts[21,64].

Overall, our results underscore the importance of mapping nonlinear and context-dependent effects, highlight the benefits of integrating these effects to improve PGS transferability, and, more broadly, pave the way for designing more inclusive and nuanced health interventions.

## Methods

### Compliance with ethical regulations and informed consent

This research has been conducted using the UK Biobank Resource under Application Number 21942, “Integrated models of complex traits in the UK Biobank” (https://www.ukbiobank.ac.uk/enable-your-research/approved-research/integrated-models-of-complex-traits-in-the-uk-biobank-the-uk-biobank). All participants of UK Biobank provided written informed consent. More information is available at https://www.ukbiobank.ac.uk/explore-your-participation/basis-of-your-participation

### GENE and Sex Interaction Score (GeneSIS)

We introduce the GENE and Sex Interaction Score (GeneSIS) methodology as an application of supervised learning for phenotypic prediction. In this initial study, we applied *L*_1_- and *L*_2_-penalized Elastic Net regression directly on the individual-level data using the batch screening iterative lasso (BASIL) algorithm implemented in the R *snpnet* package (version 2)[9,38–41,65], although other statistical and machine-learning approaches, such as statistical boosting[33,60], would be applicable as described in the discussion.

Given a phenotype vector *y* = (*y*_1_…*y*_*n*_)^T^ ∈ ℛ^*n*^ of *n* individuals and a predictor matrix *X* = (*x*_1_ … *x*_*n*_)^T^ ∈ ℛ^*n×d*^ of *n* individuals and *d* variables, we consider the following penalized generalized linear regression to fit the intercept term 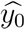 and regression coefficient vector 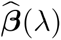 as a function of the tuning parameter λ that controls the sparsity of the solution:

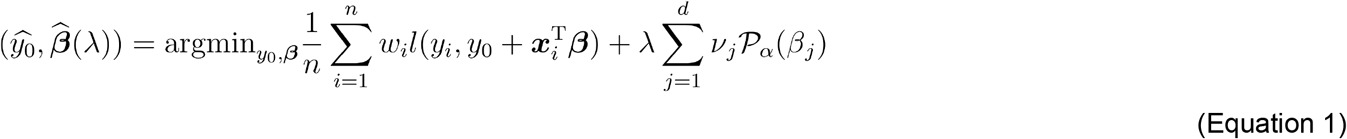

where *l*(*y*_*i*_, *η*_*i*_) is the loss contribution for the *i*-th individual and *P*_*α*_(*β*) is the Elastic Net penalization term for the coefficient *β*. With a balancing parameter between the *L*_1_-(Lasso) and *L*_2_-(Ridge) penalty, α, the Elastic Net penalization term is written as follows[40]:

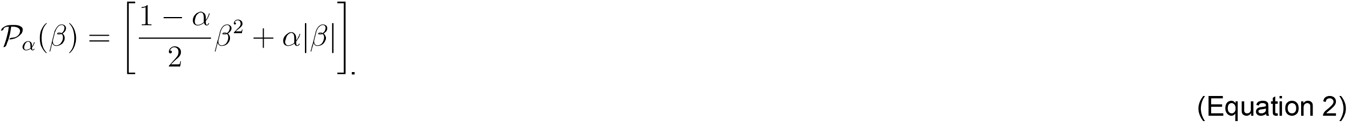

We optimize the tuning parameter, λ, based on the model’s predictive performance on the validation set and use α=0.99 (ref:[9]). As optional parameters, one may specify sample weights, *w*_*i*_, and penalty factor values, *v*_*j*_, that allow different levels of shrinkage to variables according to prior knowledge. In our application to the UK Biobank dataset, we set *v*_*j*_ =0 for covariate terms to ensure the covariate terms are unpenalized in the regression[9]. We assigned lower penalty factors for non-synonymous coding variants, as described below. One may use *w*_*i*_ =1 and *v*_*j*_ =1 as defaults.

The loss contribution for the *i*-th observation, *l*(*y*_*i*_, *η*_*i*_), depends on the types of the exponential family considered in the regression analysis of generalized linear models. For example, it is the squared loss, i.e., 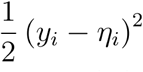, for quantitative phenotypes (Gaussian family), and it is logistic loss, i.e., −*y*_*i*_ log(*η*_*i*_) − (1 − *y*_*i*_) log(1 − *η*_*i*_), for binary phenotypes y ∈ {0, 1}^*n*^ (Binomial family). A similar model can be used for time-to-event phenotypes (Cox Proportional Hazards regression) or joint modeling of multiple phenotypes, as shown in our previous studies[38,39,45,66–68].

### GeneSIS with linear effects alone

For modeling linear effects of genetic variants[9], we consider a covariate matrix 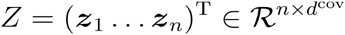 of *n* individuals and *d*^cov^ covariates and a genotype matrix 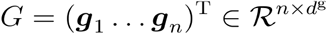 of *n* individuals and *d*^g^ variants representing the allelic count of effect allele. We use the concatenated vectors as predictors:

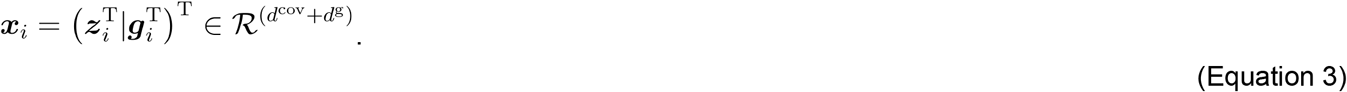

The coefficient vector is also a concatenation of two components, corresponding to covariate effects and linear effects of genetic variants, respectively:

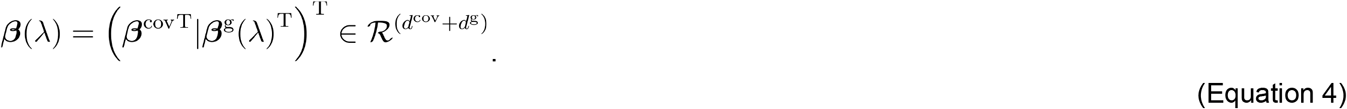

When we set covariate terms to be unpenalized (i.e., *v*_*j*_= 0 for *j* ∈ [1, …, *d*^*cov*^]),, the objective function of the penalized generalized linear regression (Equation 1) becomes the following, which is equivalent to the polygenic prediction models considered in our previous studies[9,38,45]:

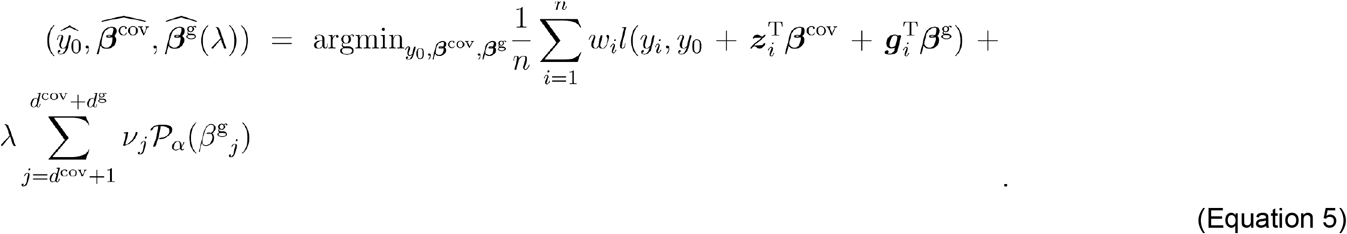

### GeneSIS with genetic dominance effects

To incorporate nonlinear genetic dominance effects, we consider a matrix, 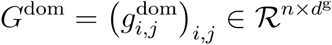, where 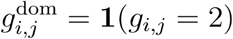 is an indicator variable representing whether the *i*-th individual is homozygous for effect allele for the *j*-th genetic variant. We use the concatenated vectors of three components as predictors:

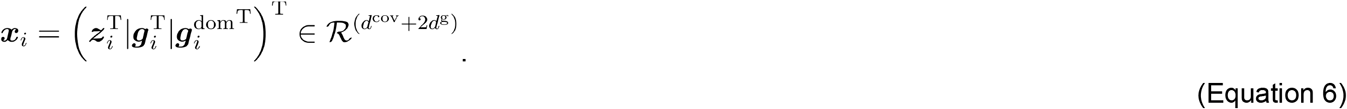

The coefficient vector now has three components:

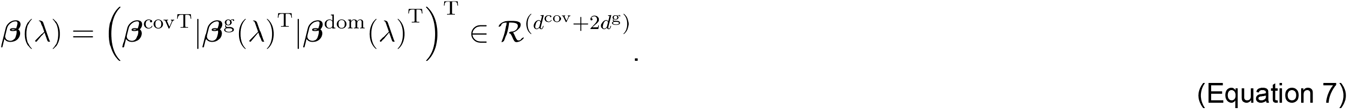

One may focus on a smaller subset of variants for genetic dominance effects instead of considering all of the *d*^g^ genetic variants.

### GeneSIS with GxS interaction effects

To incorporate gene-by-sex (GxS) interaction effects, we augment predictor variables. For example, we may consider two matrices, 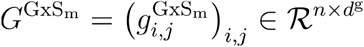 and 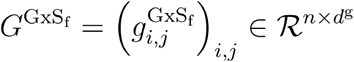, where 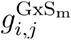 and 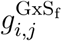 represent GxS interaction terms for male and female individuals, respectively, as follows:

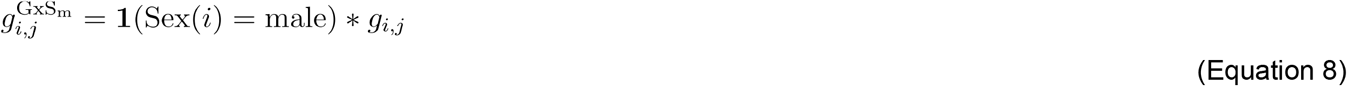

and

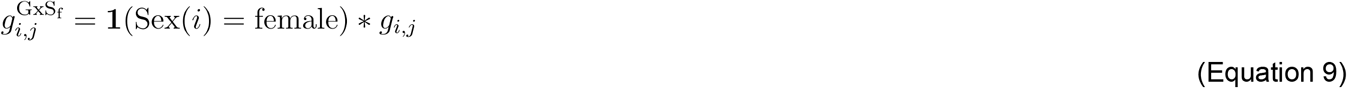

where 1(·) is an indicator function. The 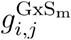 has the original genotype dosage for male individuals and the value is always set to be zero for female individuals and vice versa for 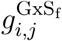. We use the concatenated vectors of four components as predictors:

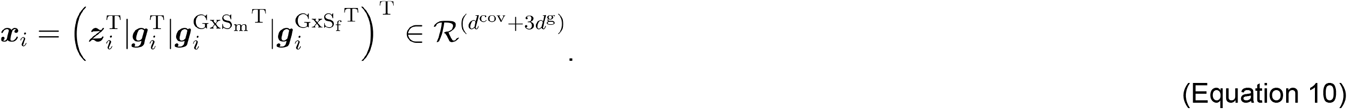

The coefficient vector now has four components:

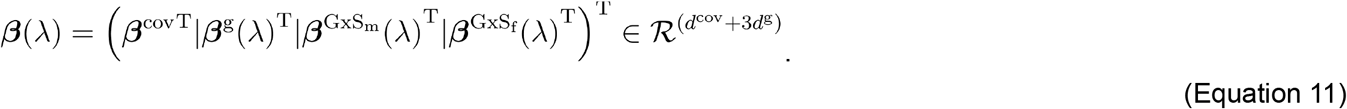

There would be other encoding of GxS interaction effects, as follows:

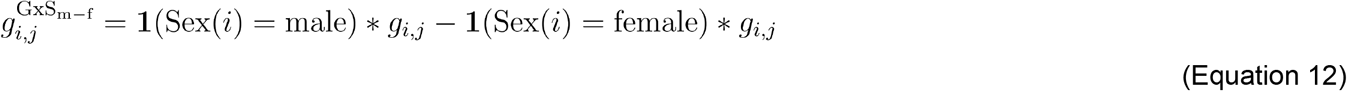

One may focus on a smaller subset of variants for the GxS interaction effect.

### GeneSIS with genetic dominance and GxS interaction effects

It is possible to consider genetic dominance and GxS interaction effects of the linear and genetic dominance effects simultaneously by using concatenated vectors of seven components as predictors:

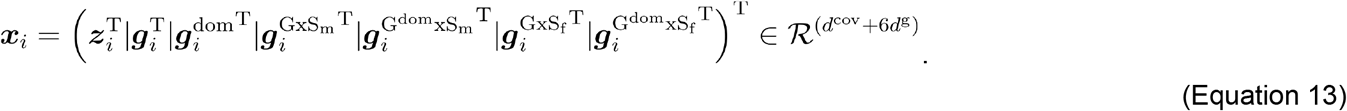

The coefficient vector now has seven components:

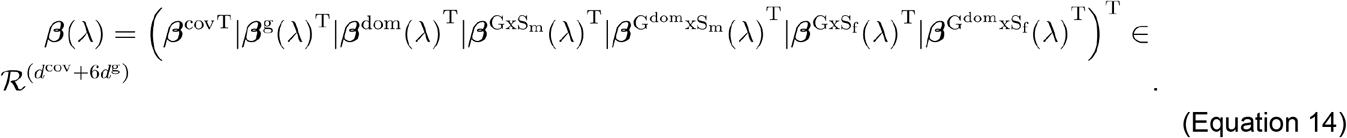

One may focus on smaller subsets of variants for genetic dominance effects and GxS interaction effects of the linear and genetic dominance effects.

### The study population in the UK Biobank resource

UK Biobank is a population-based cohort study with genomic and phenotypic datasets across about 500,000 volunteers collected across multiple sites in the United Kingdom[37,69,70]. We performed sample-level quality control and focused on n=406,659 unrelated individuals with genetic data based on the following criteria[9]: (1) used to compute principal components (UK Biobank Data Field 22020); (2) removal of sex mismatch between the sex field in the genotype dataset and phenotype sex (Data Field 31); (3) not reported in “Outliers for heterozygosity or missing rate” (Data Field 22027); (4) not reported in “Sex chromosome aneuploidy” (Data Field 22019); and (5) do not have ten or more third-degree relatives (Data Field 22021). We subsequently used a combination of self-reported ethnic background (Data Field 21000) and genetic principal components (Data Field 22009) to define four population groups (White British, non-British white, African, and South Asian) and kept the remaining unrelated individuals as Others[9,52]. We focused on individuals whose inferred biological sex is either female or male. In each population, 53.9% of unrelated individuals are female and the remaining 46.1% are male (**Table S1**). We randomly split each population group into training (70%, n=284,661), validation (10%, n=40,667), and held-out test (20%, n=81,331) sets without using phenotypes[9]. We used the training set for model fitting, the validation set for determining hyperparameters, and the held-out test sets for predictive performance evaluation. We used the same training, validation, and test sets for all tested traits[9].

### Variant annotation and quality control in UK Biobank

We used the directly genotyped dataset (release version 2), imputed genotypes (release version 3), imputed HLA allelotype (release version 2), and GRCh37 human reference genome throughout the study[37]. As in our previous study[9], we performed variant annotation with Ensembl’s Variant Effect Predictor (VEP) (version 101)[71,72] with the LOFTEE plugin[73] and ClinVar[74]. We grouped the VEP-predicted consequence of the variants into six groups[9,45]: protein-truncating variants (PTVs), protein-altering variants (PAVs), proximal coding variants (PCVs), intronic variants (Intronic), genetic variants on untranslated regions (UTR), and other non-coding variants (Others)[71,72]. For the directly genotyped dataset, we focused on variants passing the following criteria: (1) the missingness of the variant is less than 1%, considering that the two genotyping arrays (the UK BiLEVE Axiom array and UK Biobank Axiom array) cover a slightly different set of variants[37] and (2) Hardy-Weinberg disequilibrium test p-value greater than 1.0 × 10^−7^. For the imputed genotype dataset, we used the following criteria: (1) the missingness of the variant is less than 1%; (2) minor allele frequency (MAF) greater than 0.01%; (3) imputation quality score (INFO score) greater than 0.3; (4) does not present in the directly genotyped dataset; and (5) present in the HapMap Phase 3 dataset. For the HLA allelotype, we kept the imputed allelotype dosage within [0, 0.1), (0.9, 1.1), or (1.9, 2.0] and converted it to hard call[75]. We focused on the HLA allelotype with (1) missingness no more than 1% and (2) Hardy-Weinberg disequilibrium test p-value greater than 1.0 × 10^−4^. We concatenated all variants and allelotypes into one dataset using PLINK 2.0 (v2.00a3.3LM 3 Jun 2022)[76]. The quality control procedure resulted in 1,316,181 unique genetic variants and allelotypes considered in the analysis[9].

We defined the following variables to consider nonlinear genetic effects. The imputed HLA allelotypes account for complex LD structure in the major histocompatibility complex (MHC) region. We kept the imputed allelotype dosage within (1.9, 2.0] for genetic dominance effects of imputed HLA allelotypes. For GxS interaction effects, we modeled sex-specific effects in males by keeping the original genotype for male individuals and setting zero for female individuals, as in Equation 8. Lastly, we prepared variables for GxS interaction effects of genetic dominance terms for the HLA allelotypes. We dropped variables when none of the unrelated individuals considered in the analysis had non-zero values. The procedure above resulted in 2,630,335 variables considered in the analysis (**Table S3**).

### Phenotype definition in UK Biobank

In the UK Biobank resource, we focused on 99 anthropometric, blood biochemistry, blood count (hematological), and blood pressure traits (**Table S2**). Some of those phenotypes are collected at up to four instances, each of which corresponds to (1) the initial assessment visit (2006-2010), (2) the first repeat assessment visit (2012-2013), (3) the imaging visit (2014-), and (4) the first repeat imaging visit (2019-). We defined phenotype data as the median of non-missing values for each individual across the 60 quantitative traits, as described elsewhere[45,51,77].

### Genome-wide association analysis

We applied sex-stratified genome-wide association analysis with PLINK (v2.00 alpha)[76]. We computed population-specific genotype PCs for White British individuals in the UK Biobank cohort using the randomized algorithm implemented in PLINK2. We subsequently applied the GWAS analysis using age, Townsend deprivation index, array, and the top ten population-specific genotype PC loadings as covariates, using an approximation algorithm implemented as the “--glm zs omit-ref no-x-sex log10 hide-covar skip-invalid-pheno cc-residualize firth-fallback” command in PLINK2[78]. The participants of the UK Biobank cohort were genotyped on two different arrays: about 10% of participants were genotyped on the UK BiLEVE Axiom array, whereas the rest were genotyped on the UK Biobank Axiom array[37]. When genetic variants were directly measured on both arrays, we included an indicator variable “array” in the covariates, denoting whether the UK Biobank Axiom array or UK BiLEVE Axiom array was used for genotyping. We applied quantile normalization using the “--pheno-quantile-normalize” option in PLINK2. We conducted GWAS analysis using the male and female individuals separately in the White British population group.

### Sex-stratified GWAS analysis

To test the sex difference in GWAS associations, we applied the interaction effect-only test described in a recently published study[18]. Specifically, we calculated the sex-differential test statistics *T*_Diff_, applying Equation 14 shown below to association summary statistics from the sex-stratified analysis:

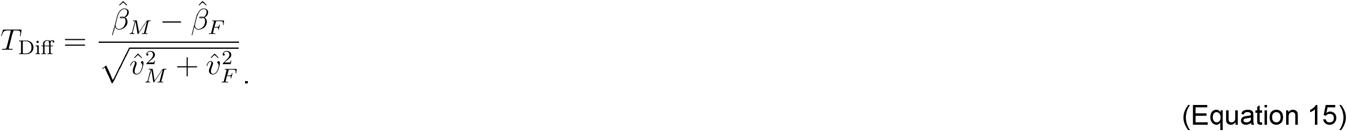

### Applying GeneSIS to the UK Biobank resource

In our application of GeneSIS to UK Biobank, we included the following variables as unpenalized covariates: age (UK Biobank Data Field 34), sex (Data Field 31), age^2^, age*sex, Townsend deprivation index (Data Field 22189), and the first 18 genotype PCs (Data Field 22009) provided by UK Biobank[37]. The genotype PCs represent genetic ancestry and account for trait mean differences associated with the genome-wide genetic ancestry. We considered linear, genetic dominance, and sex-specific GxS interaction effects in males represented in *p*=2,630,335 variables across 1,316,181 unique genetic variants and imputed allelotypes (**Table S3**). Analysis of both male- and female-specific GxS effect vectors presents technical challenges in numerical stability, partly due to the collinearity of sex-specific GxS effect vectors. In the main analysis presented in this initial study, we focused on male-specific GxS interaction effects, given that more female individuals are present in the cohort. For select traits, we incorporated both male- and female-specific GxS effect vectors and investigated the difference in predictive performance. We prioritized protein-truncating and protein-altering variants using penalty factors shown in **Table S3** to improve the interpretation of selected variants, as in our previous studies[9,45]. We imposed more penalization for nonlinear genetic effects to reduce the risk of overfitting. The specific values of penalty factors are based on heuristics[9,45], and finding the optimal values of penalty factors would be an important direction of follow-up studies as described in the Discussion.

### Comparison of linear and GxS effect size

We computed the standard deviation of the phenotype values in the training set for each trait and used it to normalize the GeneSIS effect size. We computed the first, median, and third quartile of the absolute value of normalized effect size for linear and GxS effects. We compared them across 99 traits. We fit a linear regression with a fixed intercept at zero and reported the slope as an aggregated measure of the difference in the magnitude of effect size between linear and GxS effects across traits.

### PGS performance evaluation and model selection

We evaluated the predictive performance (*R*^2^) of (1) PGS (non-covariate-only) models, (2) covariate-only models, and (3) full models that considered both covariates and genotypes using the held-out test set (**Table S2**-**S3**). Unless indicated otherwise, we reported the full models’ predictive performance in the remainder of the main text. We evaluated the 95% confidence interval of predictive performance using the approximate standard error of the *R*^2^ metric[79,80].

In our application of GeneSIS to UK Biobank, we used the predictive performance of the linear-only iPGS model as the baseline[9]. We assessed the significance of the difference in *R*^2^ between the GeneSIS model and the iPGS model using the delta method implemented in the r2redux package in R[79,81].

We performed model selection between GeneSIS and linear-only iPGS for each (trait, population group) pair using the predictive performance in the validation set. We selected the GeneSIS model if GeneSIS showed improved performance with statistical significance of nominal p-values less than 0.05. We selected iPGS models otherwise (**Fig S3**).

### Predictive performance evaluations reported in the PGS catalog

To compare the predictive performance of the GeneSIS model against that of publicly available PGS evaluations in the PGS catalog[44], we evaluated the predictive performance of the GeneSIS model using partial-*R*^2^ adjusted for covariate effects using Equation 16:

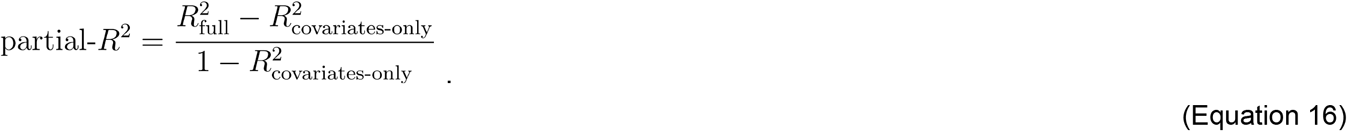

We used the 95% confidence intervals of the original *R*^2^ _full_ (Equation 16) to compute the 95% confidence intervals of partial-*R*^2^. We found four studies with publicly available PGS evaluations for hip circumference for African-ancestry individuals (https://www.pgscatalog.org/trait/EFO_0005093/)[9,44–47]. We converted partial correlation (partial-r) to partial-*R*^2^ by squaring its value.

### Locus views of select genetic variants

We used igv-notebook Python package to visualize genome-browser view of a few example genetic variants selected from the GeneSIS model[82].

### Phenome-wide association analysis

We investigated the phenome-wide association of genetic variants with GxS interaction effects in the GeneSIS models using the Global Biobank Engine[51]. We obtained the association summary statistics for the selected genetic variants based on their coordinates on the GRCh37 reference genome. For example, the association profile for rs1260326 (a missense variant in *GCKR*) is from the following webpage: https://biobankengine.stanford.edu/RIVAS_HG19/variant/2-27730940-T-C. We subsequently focused on associations with nominal p-value < 1 × 10^−10^ across traits in the following categories: anthropometry, arterial stiffness, biomarkers, blood assays, blood pressure, and sex-specific factors[51,52,77].

### Enrichment to biological processes

To investigate the relevant biological processes in the nonlinear genetic effects captured in GeneSIS models, we applied the Genomic Regions Enrichment of Annotations Tool (GREAT) (version 4.0.4)[42,43]. To evaluate the enriched biological processes captured in linear and nonlinear effects in GeneSIS models, we took the top 1000 genetic variants according to the effect size and applied the GREAT enrichment against the default genome-wide background. To evaluate the enrichment of genetic variants with GxS interaction effects, we took the variants with non-zero GxS interaction effects as the foreground set and evaluated their relative enrichment against all the variants included in the GeneSIS model (background set). In both analyses, we filtered ontology terms based on the number of genes annotated for the term, focusing on terms with 5 to 500 annotated genes.

### Statistics

For computational and statistical analysis, we used Jupyter Notebook[83], R[84], R Tidyverse package[85], and GNU parallel[86] (https://www.gnu.org/software/parallel/). For visualization, we used ggplot2[87] with ggrepel[88] (https://github.com/slowkow/ggrepel) and ggrastr[89] (https://CRAN.R-project.org/package=ggrastr) packages. The p-values were computed from two-sided tests unless otherwise specified.

## Data and Code Availability

The analyses presented in this study were based on the individual-level data accessed through UK Biobank: https://www.ukbiobank.ac.uk. For the PGS analyses with GeneSIS and linear-only inclusive PGS, we used the BASIL algorithm implemented in the R *snpnet* package version 2 (https://github.com/rivas-lab/snpnet/tree/compact). The igv-notebook is available at https://github.com/igvteam/igv-notebook. The GREAT enrichment web tool (version 4) is available at https://great.stanford.edu/.

## Acknowledgments

This work was supported in part by the National Institutes of Health grants AG054012, AG058002, MH109978, AG062377, AG081017, NS129032, AG077227, NS110453, NS115064, AG062335, AG074003, NS127187, AG067151, MH119509, HG008155, and DA053631 (M.K.). We thank William F Li, Patricia Purcell, Amy Grayson, and the members of the Kellis lab for their scientific suggestions and feedback on earlier versions of the manuscript. The content is solely the responsibility of the authors and does not necessarily represent the official views of the funding agencies; funders had no role in study design, data collection, data analysis, the decision to publish, or the preparation of the manuscript.

## Author Contributions

Y.T. conceived and designed the study; Y.T. developed the computational framework and conducted data analysis; Y.T. and M.K. interpreted the results; M.K. was responsible for funding acquisition; and Y.T. wrote the manuscript with feedback from M.K.

